## Supplemental Document 1 for "Vertex protein PduN tunes encapsulated pathway performance by dictating bacterial metabolosome morphology"

Mills *et al.*

### Supplemental Methods: Calculation of Bending Potential

The bending potential between the hexamer and pentamer, $V(\theta_B)$, is calculated from the forces on the pentamer at a given bending angle, according to the definition of the force, $F_{\theta_B}$:

$$F_{\theta_B} = -\frac{\partial V}{\partial \theta_B}$$

Here *V* is the general interaction potential between the hexamer and pentamer, so it is necessary to measure only the forces in the $\theta_B$-direction. Using the GROMACS simulation engine [1], we constrain the proteins in Cartesian directions but not the angle itself. Instead, we constrain the backbone of the hexamer to not move in any cartesian direction while the pentamer is allowed to move in the yz-plane. We then measure the forces in the z-direction, $\frac{-\partial V}{dz}$, as logged from the simulations, $F_{gromacs}$, and convert to the force in the $\theta_B$-direction as follows:

$$F_{\theta_B}(z) = \frac{-\partial V}{\partial \theta_B} = \frac{-\partial V}{\partial z}\frac{\partial z}{\partial \theta_B} = \frac{-\partial V}{\partial z} cos\theta_B = -F_Z cos\theta_B = -F_{gromacs} cos\theta_B$$

In the simulations, components of $F_Z$ not in the $\theta_B$-direction are cancelled out by the constraints on the hexamer and thus all net forces and movement of the protein are in the $\theta_B$-direction. Since we can measure the bending angle, $\theta_B$, in the simulation, we can create a one-to-one map between *z* and $\theta_B$, where *z* is the distance between the centers of mass of the pentamer and hexamer (Supplemental Figure S6).

We run simulations at different angles and are thus able to calculate a potential of mean force by performing a discrete summation over those angles.

$$V(\theta_{B,n}(z_n)) - V(\theta_{B,0}(z_0)) = -\sum_{i=0}^{n-1} \langle F_{\theta_B}(\langle z_i \rangle)\rangle (z_{i+1} - z_i)$$

We are careful to use many different “windows” (i.e. make n large and $(z_{i+1} - z_i)$ small) to calculate the mean forces at mean positions in a pseudo-continuous manner with overlap between states, especially near the minimum. In Supplemental Figure S7, each color represents one “window.” The same method applies to the case of two hexamers.

A general version of this method is used to calculate the total interaction strength be computing the potential of mean force as a function of the distance, *R*, between the proteins until they are no longer interacting at $R_0$.

$$V(R_n) - V(R_0) = -\sum_{i=0}^{n-1} \langle F(\langle R_i \rangle)\rangle (R_{i+1} - R_i)$$

Here, $F(R)$ is the force calculated from simulation and does not require adjustments for angular components.

**Supplemental Discussion 1: Systems-level kinetic model**

*Discrepancies between model and experimental data*

In the main text, we noted that there were two main discrepancies between the systems-level kinetic model and our experimental data: (1) propionate and 1-propanol are eventually consumed in our experiments, and (2) absolute propionaldehyde concentrations observed differ from those predicted in the model. To point (1), this discrepancy is expected, as there are no terms in our model that account for the uptake of propionate into central metabolism, which is what occurs in our experiments. Development of the model to accurately capture this phenomenon is certainly of interest for future study, but does not strongly impact the differences we wish to explore here, which are the changes in propionaldehyde buildup in our different compartment geometries. We can be confident that down-stream reactions have low impact on propionaldehyde concentrations because of our sensitivity analysis. The main way in which we expect that excess propionate and 1-propanol would affect propionaldehyde is through the reverse reactions. However, varying the rate of the reverse reaction in the sensitivity analysis has negligible effect on peak propionaldehyde concentration (Supplemental Figures S4 and S5). So, we expect that including propionate and 1-propanol consumption in the model would have negligible effect on the propionaldehyde peak. To point (2), it has been noted in previous growth studies that the observed propionaldehyde level in the media can be impacted by a suite of growth factors (temperature, shaking speed), due to the volatility of propionaldehyde [2]. It is thus reasonable to expect that the observed propionaldehyde level in the media may be lower than predicted by our model.

*Calculating the number of Pdu MTs assuming either volume or surface area matches that of Pdu MCPs*

We noted in the main text that we compared the base MCP case to the MT case by keeping as many parameters the same in the MT model, and that this included total enzyme number per cell. We then considered two limiting cases—one in which total compartment surface area is the same in both MCP and MT models (resulting in an increased enzyme concentration in MTs), and a second in which total volume is the same in both MCP and MT models, which keeps the enzyme concentration the same in both geometries, but results in increased surface area in the MTs. We calculated the number of MTs per cell in these two limiting cases. First, if MCP surface area is conserved (*i.e.*, all shell proteins that make up the shell of 15 MCPs will contribute to MT surface area), then there would be 2.4 Pdu MTs per cell containing 54% of the original MCP volume. Alternatively, if MCP internal volume is conserved (*i.e.*, all enzymes inside 15 MCPs are now inside MTs, and requisite shell protein is assumed available), then there would be 4.4 MTs per cell encased in 1.9 times the original MCP surface area.

*Sensitivity Analysis of Systems-Level Model*

We used a local sensitivity analysis to identify enzyme and compartment features that had the highest impact on propionaldehyde buildup in the two cases (1) spherical MCPs and (2) Pdu MTs with increased surface area (the constant volume case described in the main text). We varied parameters by 10% and observed the resulting maximum propionaldehyde concentration for both models. This analysis revealed that there were two common features in both geometries that determined propionaldehyde buildup outside of the cell: (1) the kinetics of the PduP and PduQ enzymes, and (2) the overall transport of substrates in and out of the compartment volume (Supplemental Figures S4 and S5). The decrease in propionaldehyde buildup with enhanced PduP/PduQ activity (either by increasing $V_{max}$ or by decreasing $K_M$ for the forward reaction) is expected, as PduP and PduQ both have relatively slow kinetics compared to PduCDE, which rapidly generates propionaldehyde (resulting in buildup) [3], [4], [5], [6]. The main compartment features that influence peak propionaldehyde level are compartment surface area and permeability—as the compartment surface area or permeability increases, the access of PduCDE to the 1,2-propandiol substrate increases, resulting in more rapid and thus substantial buildup of the toxic propionaldehyde intermediate. Interestingly, in the Pdu MT constant volume case where there is increased surface area (Supplemental Figure S5), the peak propionaldehyde level is also sensitive to the $V_{max}$ (forward) of the PduCDE enzyme. This is distinct from the spherical MCP case, where the kinetics of the PduCDE enzyme had little effect on peak propionaldehyde level. This is because the increased surface area that the Pdu MTs have in the constant volume case (1.9 times higher than that of spherical MCPs) increases the overall flux of 1,2-propanediol into the compartment volume, shifting the system into a regime where the rate of the reaction mediated by the PduCDE enzyme is similar to the rate at which 1,2-propanediol enters the compartment. In other words, higher surface area Pdu MTs are no longer rate limiting and the reaction mediated by the PduCDE enzyme becomes rate limiting.

**Supplemental Discussion 2: Glycine 52 point mutant to asparagine**

We noted in the main text that unlike other PduN point mutants at residue G52, PduN-G52N has a mixed population of MCP and MT-like structures. These results suggest that the PduN-G52N is able to facilitate Pdu MCP vertex capping to some extent. This finding was initially surprising, as the asparagine side chain is more dissimilar to the native glycine residue than a smaller point mutant like alanine. However, a more detailed look at the crystal structure of the pentamer-hexamer interface in the crystal structure of homologous shell proteins reveals that the backbone carbonyl in the G52 equivalent residue participates in a hydrogen bond with a lysine on the hexamer, suggesting that this hydrogen bond formation contributes to the stability of this interface [7], [8]. We thus hypothesize that the amidic carbonyl group in the asparagine side chain contributes to a similar hydrogen bonding interaction [9] that partially compensates for the steric penalty associated with inserting a large side chain in this position.

**SupplementalTable S1.** Strains used in this study

| **Strain** | **Organism** | **Genotype** |
|---|---|---|
| DTE003 | *S. enterica* serovar Typhimurium LT2 | Wild type |
| CEMS028 | *S. enterica* serovar Typhimurium LT2 | Δ*pduN* |
| TMDS025 | *S. enterica* serovar Typhimurium LT2 | Δ*pduD::ssD-GFPmut2* |
| CEMS055 | *S. enterica* serovar Typhimurium LT2 | Δ*pduD::ssD-GFPmut2* Δ*pduN* |
| CMJS256 | *S. enterica* serovar Typhimurium LT2 | Δ*pocR* |
| NWKs083 | *S. enterica* serovar Typhimurium LT2 | Δ*pduA* Δ*pduJ* |
| CEMS234 | *S. enterica* serovar Typhimurium LT2 | Δ*pduD::ssD-GFPmut2* Δ*pduN::pduN-G52A* |
| CEMS235 | *S. enterica* serovar Typhimurium LT2 | Δ*pduD::ssD-GFPmut2* Δ*pduN::pduN-G52L* |
| CEMS236 | *S. enterica* serovar Typhimurium LT2 | Δ*pduD::ssD-GFPmut2* Δ*pduN::pduN-G52M* |
| CEMS237 | *S. enterica* serovar Typhimurium LT2 | Δ*pduD::ssD-GFPmut2* Δ*pduN::pduN-G52N* |
| CEMS238 | *S. enterica* serovar Typhimurium LT2 | Δ*pduD::ssD-GFPmut2* Δ*pduN::pduN-G52P* |
| CEMS239 | *S. enterica* serovar Typhimurium LT2 | Δ*pduD::ssD-GFPmut2* Δ*pduN::pduN-G52C* |
| CEMS240 | *S. enterica* serovar Typhimurium LT2 | Δ*pduD::ssD-GFPmut2* Δ*pduN::pduN-G52E* |
| CEMS241 | *S. enterica* serovar Typhimurium LT2 | Δ*pduD::ssD-GFPmut2* Δ*pduN::pduN-G52H* |
| CEMS242 | *S. enterica* serovar Typhimurium LT2 | Δ*pduD::ssD-GFPmut2* Δ*pduN::pduN-G52I* |
| CEMS243 | *S. enterica* serovar Typhimurium LT2 | Δ*pduD::ssD-GFPmut2* Δ*pduN::pduN-G52K* |

| | | |
|---|---|---|
| CEMS244 | *S. enterica* serovar Typhimurium LT2 | Δ*pduD::ssD-GFPmut2* Δ*pduN::pduN-G52T* |
| CEMS245 | *S. enterica* serovar Typhimurium LT2 | Δ*pduD::ssD-GFPmut2* Δ*pduN::pduN-G52W* |
| CEMS246 | *S. enterica* serovar Typhimurium LT2 | Δ*pduD::ssD-GFPmut2* Δ*pduN::pduN-G52D* |
| CEMS247 | *S. enterica* serovar Typhimurium LT2 | Δ*pduD::ssD-GFPmut2* Δ*pduN::pduN-G52F* |
| CEMS248 | *S. enterica* serovar Typhimurium LT2 | Δ*pduD::ssD-GFPmut2* Δ*pduN::pduN-G52Q* |
| CEMS249 | *S. enterica* serovar Typhimurium LT2 | Δ*pduD::ssD-GFPmut2* Δ*pduN::pduN-G52V* |
| CEMS250 | *S. enterica* serovar Typhimurium LT2 | Δ*pduD::ssD-GFPmut2* Δ*pduN::pduN-G52Y* |
| CEMS251 | *S. enterica* serovar Typhimurium LT2 | Δ*pduD::ssD-GFPmut2* Δ*pduN::pduN-G52R* |
| CEMS252 | *S. enterica* serovar Typhimurium LT2 | Δ*pduD::ssD-GFPmut2* Δ*pduN::pduN-G52S* |
| CEMS253 | *S. enterica* serovar Typhimurium LT2 | Δ*pduD::ssD-GFPmut2* Δ*pduN::pduN-T88A* |
| CEMS254 | *S. enterica* serovar Typhimurium LT2 | Δ*pduD::ssD-GFPmut2* Δ*pduN::pduN-T88L* |
| CEMS255 | *S. enterica* serovar Typhimurium LT2 | Δ*pduD::ssD-GFPmut2* Δ*pduN::pduN-T88M* |
| CEMS256 | *S. enterica* serovar Typhimurium LT2 | Δ*pduD::ssD-GFPmut2* Δ*pduN::pduN-T88F* |
| CEMS257 | *S. enterica* serovar Typhimurium LT2 | Δ*pduD::ssD-GFPmut2* Δ*pduN::pduN-T88W* |
| CEMS258 | *S. enterica* serovar Typhimurium LT2 | Δ*pduD::ssD-GFPmut2* Δ*pduN::pduN-T88K* |
| CEMS259 | *S. enterica* serovar Typhimurium LT2 | Δ*pduD::ssD-GFPmut2* Δ*pduN::pduN-T88G* |

| | | |
|---|---|---|
| CEMS260 | *S. enterica* serovar Typhimurium LT2 | Δ*pduD::ssD-GFPmut2* Δ*pduN::pduN-T88Q* |
| CEMS261 | *S. enterica* serovar Typhimurium LT2 | Δ*pduD::ssD-GFPmut2* Δ*pduN::pduN-T88E* |
| CEMS262 | *S. enterica* serovar Typhimurium LT2 | Δ*pduD::ssD-GFPmut2* Δ*pduN::pduN-T88P* |
| CEMS263 | *S. enterica* serovar Typhimurium LT2 | Δ*pduD::ssD-GFPmut2* Δ*pduN::pduN-T88V* |
| CEMS264 | *S. enterica* serovar Typhimurium LT2 | Δ*pduD::ssD-GFPmut2* Δ*pduN::pduN-T88I* |
| CEMS265 | *S. enterica* serovar Typhimurium LT2 | Δ*pduD::ssD-GFPmut2* Δ*pduN::pduN-T88S* |
| CEMS266 | *S. enterica* serovar Typhimurium LT2 | Δ*pduD::ssD-GFPmut2* Δ*pduN::pduN-T88Y* |
| CEMS267 | *S. enterica* serovar Typhimurium LT2 | Δ*pduD::ssD-GFPmut2* Δ*pduN::pduN-T88H* |
| CEMS268 | *S. enterica* serovar Typhimurium LT2 | Δ*pduD::ssD-GFPmut2* Δ*pduN::pduN-T88R* |
| CEMS269 | *S. enterica* serovar Typhimurium LT2 | Δ*pduD::ssD-GFPmut2* Δ*pduN::pduN-T88N* |
| CEMS270 | *S. enterica* serovar Typhimurium LT2 | Δ*pduD::ssD-GFPmut2* Δ*pduN::pduN-T88D* |
| CEMS271 | *S. enterica* serovar Typhimurium LT2 | Δ*pduD::ssD-GFPmut2* Δ*pduN::pduN-T88C* |
| CEMS278 | *S. enterica* serovar Typhimurium LT2 | Δ*pduN::pduN-G52C* |
| CEMS279 | *S. enterica* serovar Typhimurium LT2 | Δ*pduN::pduN-G52N* |
| CEMS280 | *S. enterica* serovar Typhimurium LT2 | Δ*pduN::pduN-T88A* |
| TUC01 [10], [11] | *E. coli* W3110 | *gal490 pglΔ8 λcI857 Δ(cro-bioA) int<>cat/sacB* |

**SupplementalTable S2.** Plasmids used in this study

| Name | Plasmid | Origin | Resistance |
|---|---|---|---|
| CMJ144 | pBAD33t-PduN-FLAG | p15A | Chloramphenicol |
| pCEM008 | pBAD33t-PduN-6xHis | p15A | Chloramphenicol |
| pCEM027 | pBAD33t-PduN1-79-ggsfGFP-6xHis | p15A | Chloramphenicol |
| pCEM030 | pBAD33t-PduN-T88A-6xHis | p15A | Chloramphenicol |
| pCEM031 | pBAD33t-PduN-T88L-6xHis | p15A | Chloramphenicol |
| pCEM032 | pBAD33t-PduN-T88M-6xHis | p15A | Chloramphenicol |
| pCEM033 | pBAD33t-PduN-T88F-6xHis | p15A | Chloramphenicol |
| pCEM034 | pBAD33t-PduN-T88W-6xHis | p15A | Chloramphenicol |
| pCEM035 | pBAD33t-PduN-T88K-6xHis | p15A | Chloramphenicol |
| pCEM036 | pBAD33t-PduN-T88G-6xHis | p15A | Chloramphenicol |
| pCEM037 | pBAD33t-PduN-T88Q-6xHis | p15A | Chloramphenicol |
| pCEM038 | pBAD33t-PduN-T88E-6xHis | p15A | Chloramphenicol |
| pCEM039 | pBAD33t-PduN-T88P-6xHis | p15A | Chloramphenicol |
| pCEM040 | pBAD33t-PduN-T88V-6xHis | p15A | Chloramphenicol |
| pCEM041 | pBAD33t-PduN-T88I-6xHis | p15A | Chloramphenicol |
| pCEM042 | pBAD33t-PduN-T88S-6xHis | p15A | Chloramphenicol |
| pCEM043 | pBAD33t-PduN-T88Y-6xHis | p15A | Chloramphenicol |
| pCEM044 | pBAD33t-PduN-T88H-6xHis | p15A | Chloramphenicol |
| pCEM045 | pBAD33t-PduN-T88R-6xHis | p15A | Chloramphenicol |
| pCEM046 | pBAD33t-PduN-T88N-6xHis | p15A | Chloramphenicol |
| pCEM047 | pBAD33t-PduN-T88D-6xHis | p15A | Chloramphenicol |
| pCEM048 | pBAD33t-PduN-T88C-6xHis | p15A | Chloramphenicol |

| | | | |
|---|---|---|---|
| pADJ001 | pBAD33t-PduN-G52A-6xHis | p15A | Chloramphenicol |
| pADJ002 | pBAD33t-PduN-G52L-6xHis | p15A | Chloramphenicol |
| pADJ003 | pBAD33t-PduN-G52M-6xHis | p15A | Chloramphenicol |
| pADJ004 | pBAD33t-PduN-G52N-6xHis | p15A | Chloramphenicol |
| pADJ005 | pBAD33t-PduN-G52P-6xHis | p15A | Chloramphenicol |
| pADJ006 | pBAD33t-PduN-G52C-6xHis | p15A | Chloramphenicol |
| pADJ007 | pBAD33t-PduN-G52E-6xHis | p15A | Chloramphenicol |
| pADJ008 | pBAD33t-PduN-G52H-6xHis | p15A | Chloramphenicol |
| pADJ009 | pBAD33t-PduN-G52I-6xHis | p15A | Chloramphenicol |
| pADJ010 | pBAD33t-PduN-G52K-6xHis | p15A | Chloramphenicol |
| pADJ011 | pBAD33t-PduN-G52T-6xHis | p15A | Chloramphenicol |
| pADJ012 | pBAD33t-PduN-G52W-6xHis | p15A | Chloramphenicol |
| pADJ013 | pBAD33t-PduN-G52D-6xHis | p15A | Chloramphenicol |
| pADJ014 | pBAD33t-PduN-G52F-6xHis | p15A | Chloramphenicol |
| pADJ015 | pBAD33t-PduN-G52Q-6xHis | p15A | Chloramphenicol |
| pADJ016 | pBAD33t-PduN-G52V-6xHis | p15A | Chloramphenicol |
| pADJ017 | pBAD33t-PduN-G52Y-6xHis | p15A | Chloramphenicol |
| pADJ018 | pBAD33t-PduN-G52R-6xHis | p15A | Chloramphenicol |
| pADJ019 | pBAD33t-PduN-G52S-6xHis | p15A | Chloramphenicol |
| pSIM6 [11] | λ Red system repressed by cI857 | pSC101 $repA^{ts}$ | Ampicillin |

**SupplementalTable S3.** Primers used in this study.

| Name | Purpose | Description | Sequence (5' → 3') |
|---|---|---|---|
| oCEM 020 | Recombineering | Amplify *cat/sacB* with homology upstream of *pduN* For | taattaagcaggagtaaatcatgcatctggcacgagtcacTGTGACGGAAGATCACTTCG |
| oCEM 014 | Recombineering | Amplify *cat/sacB* with homology downstream of *pduN* Rev | agcgtcacctgttcgggtataaatcgccataaccgccccttaacacgaaaATCAAAGGGAAAACTGTCCATAT |
| oCEM 095 | Recombineering | Amplify *pduN* point mutants with homology upstream of *pduN* For | caatgcgcggaatattcaattaattaagcaggagtaaatcATGCATCTGGCACGAG |
| oCEM 096 | Recombineering | Amplify *pduN-G52* point mutants with homology downstream of *pduN* Rev | cagcgtcacctgttcgggtataaatcgccataaccgccccTTAACACGAAAGCGTATCTACAAT |
| oCEM 097 | Recombineering | Amplify *pduN-T88A* point mutant with homology downstream of *pduN* Rev | cagcgtcacctgttcgggtataaatcgccataaccgccccTTAACACGAAAG**<u>CGC</u>**ATCTACAAT |
| oCEM 098 | Recombineering | Amplify *pduN-T88L* point mutant with homology downstream of *pduN* Rev | cagcgtcacctgttcgggtataaatcgccataaccgccccTTAACACGAAAG**<u>CAG</u>**ATCTACAAT |
| oCEM 099 | Recombineering | Amplify *pduN-T88M* point mutant with homology downstream of *pduN* Rev | cagcgtcacctgttcgggtataaatcgccataaccgccccTTAACACGAAAG**<u>CAT</u>**ATCTACAAT |
| oCEM 100 | Recombineering | Amplify *pduN-T88F* point mutant with homology downstream of *pduN* Rev | cagcgtcacctgttcgggtataaatcgccataaccgccccTTAACACGAAAG**<u>AAA</u>**ATCTACAAT |

| | | | |
|---|---|---|---|
| oCEM 101 | Recombineering | Amplify *pduN-T88W* point mutant with homology downstream of *pduN* Rev | cagcgtcacctgttcgggtataaatcgccataaccgccccTTAACACGAAAG**CCA**ATCTACAAT |
| oCEM 102 | Recombineering | Amplify *pduN-T88K* point mutant with homology downstream of *pduN* Rev | cagcgtcacctgttcgggtataaatcgccataaccgccccTTAACACGAAAG**TTT**ATCTACAAT |
| oCEM 103 | Recombineering | Amplify *pduN-T88G* point mutant with homology downstream of *pduN* Rev | cagcgtcacctgttcgggtataaatcgccataaccgccccTTAACACGAAAG**GCC**ATCTACAAT |
| oCEM 104 | Recombineering | Amplify *pduN-T88Q* point mutant with homology downstream of *pduN* Rev | cagcgtcacctgttcgggtataaatcgccataaccgccccTTAACACGAAAG**CTG**ATCTACAAT |
| oCEM 105 | Recombineering | Amplify *pduN-T88E* point mutant with homology downstream of *pduN* Rev | cagcgtcacctgttcgggtataaatcgccataaccgccccTTAACACGAAAG**TTC**ATCTACAAT |
| oCEM 106 | Recombineering | Amplify *pduN-T88P* point mutant with homology downstream of *pduN* Rev | cagcgtcacctgttcgggtataaatcgccataaccgccccTTAACACGAAAG**CGG**ATCTACAAT |
| oCEM 107 | Recombineering | Amplify *pduN-T88V* point mutant with homology downstream of *pduN* Rev | cagcgtcacctgttcgggtataaatcgccataaccgccccTTAACACGAAAG**CAC**ATCTACAAT |
| oCEM 108 | Recombineering | Amplify *pduN-T88I* point mutant with homology downstream of *pduN* Rev | cagcgtcacctgttcgggtataaatcgccataaccgccccTTAACACGAAAG**AAT**ATCTACAAT |

| | | | |
|---|---|---|---|
| oCEM 109 | Recombineering | Amplify *pduN-T88S* point mutant with homology downstream of *pduN* Rev | cagcgtcacctgttcgggtataaatcgccata accgccccTTAACACGAAAG**GCT**A TCTACAAT |
| oCEM 110 | Recombineering | Amplify *pduN-T88Y* point mutant with homology downstream of *pduN* Rev | cagcgtcacctgttcgggtataaatcgccata accgccccTTAACACGAAAG**ATA**A TCTACAAT |
| oCEM 111 | Recombineering | Amplify *pduN-T88H* point mutant with homology downstream of *pduN* Rev | cagcgtcacctgttcgggtataaatcgccata accgccccTTAACACGAAAG**ATG**A TCTACAAT |
| oCEM 112 | Recombineering | Amplify *pduN-T88R* point mutant with homology downstream of *pduN* Rev | cagcgtcacctgttcgggtataaatcgccata accgccccTTAACACGAAAG**GCG**A TCTACAAT |
| oCEM 113 | Recombineering | Amplify *pduN-T88N* point mutant with homology downstream of *pduN* Rev | cagcgtcacctgttcgggtataaatcgccata accgccccTTAACACGAAAG**GTT**A TCTACAAT |
| oCEM 114 | Recombineering | Amplify *pduN-T88D* point mutant with homology downstream of *pduN* Rev | cagcgtcacctgttcgggtataaatcgccata accgccccTTAACACGAAAG**ATC**A TCTACAAT |
| oCEM 115 | Recombineering | Amplify *pduN-T88C* point mutant with homology downstream of *pduN* Rev | cagcgtcacctgttcgggtataaatcgccata accgccccTTAACACGAAAG**GCA**A TCTACAAT |
| oCEM 015 | Recombineering | Knockout *pduN* | ATGCATCTGGCACGAGTCACGG GCGCGGTTtttcgtgttaaggggcggttat ggcgattt |

| | | | |
|---|---|---|---|
| oCEM 021 | Sequencing | Amplify from upstream of *pduN* | gcagcgcattgtcgaggagattgt |
| oCEM 022 | Sequencing | Amplify from downstream of *pduN* | ctgctggatggcctcgagtaaggt |
| oCEM 023 | Sequencing | Amplify from upstream of *pduN* | ttcggtgctgagctatcgcgatgt |
| oCEM 024 | Sequencing | Amplify from downstream of *pduN* | cctcgagtaaggtgcggtggtttt |
| oCEM 038 | Gibson cloning | Amplify pBAD33t-PduN1-79 backbone with homology to constitutively active sfGFP dropout marker For | tgaggtctcaCATCACCATCACCATCAC |
| oCEM 039 | Gibson cloning | Amplify pBAD33t-PduN1-79 backbone with homology to constitutively active sfGFP dropout marker Rev | ggaggtctcaGTCAATGGCCTCATTTGG |
| oCEM 040 | Gibson cloning | Amplify constitutively active sfGFP with homology to pBAD33t-PduN1-79 backbone For | ggccattgacTGAGACCTCCCTATCAGTG |
| oCEM 041 | Gibson cloning | Amplify constitutively active sfGFP with homology to pBAD33t-PduN1-79 backbone Rev | gatggtgatgTGAGACCTCATTTGTACAG |
| oCEM 043 | Golden Gate cloning | Amplify single-stranded DNA with PduN point mutant For | gggaGGTCTCaTGAC |
| oCEM 044 | Golden Gate cloning | Amplify single-stranded DNA with PduN point mutant Rev | cccaGGTCTCaGATG |

| | | | |
|---|---|---|---|
| oCEM 045 | Golden Gate cloning | PduN-80-91-His-T88A oligo | aGGTCTCaTGACCTCGCCGTTGTCGGCATTGTAGAT**GCG**CTTTCGTGTCATCtGAGACCt |
| oCEM 046 | Golden Gate cloning | PduN-80-91-His-T88L oligo | aGGTCTCaTGACCTCGCCGTTGTCGGCATTGTAGAT**CTG**CTTTCGTGTCATCtGAGACCt |
| oCEM 047 | Golden Gate cloning | PduN-80-91-His-T88M oligo | aGGTCTCaTGACCTCGCCGTTGTCGGCATTGTAGAT**ATG**CTTTCGTGTCATCtGAGACCt |
| oCEM 048 | Golden Gate cloning | PduN-80-91-His-T88F oligo | aGGTCTCaTGACCTCGCCGTTGTCGGCATTGTAGAT**TTT**CTTTCGTGTCATCtGAGACCt |
| oCEM 049 | Golden Gate cloning | PduN-80-91-His-T88W oligo | aGGTCTCaTGACCTCGCCGTTGTCGGCATTGTAGAT**TGG**CTTTCGTGTCATCtGAGACCt |
| oCEM 050 | Golden Gate cloning | PduN-80-91-His-T88K oligo | aGGTCTCaTGACCTCGCCGTTGTCGGCATTGTAGAT**AAA**CTTTCGTGTCATCtGAGACCt |
| oCEM 051 | Golden Gate cloning | PduN-80-91-His-T88G oligo | aGGTCTCaTGACCTCGCCGTTGTCGGCATTGTAGAT**GGC**CTTTCGTGTCATCtGAGACCt |
| oCEM 052 | Golden Gate cloning | PduN-80-91-His-T88Q oligo | aGGTCTCaTGACCTCGCCGTTGTCGGCATTGTAGAT**CAG**CTTTCGTGTCATCtGAGACCt |
| oCEM 053 | Golden Gate cloning | PduN-80-91-His-T88E oligo | aGGTCTCaTGACCTCGCCGTTGTCGGCATTGTAGAT**GAA**CTTTCGTGTCATCtGAGACCt |
| oCEM 054 | Golden Gate cloning | PduN-80-91-His-T88P oligo | aGGTCTCaTGACCTCGCCGTTGTCGGCATTGTAGAT**CCG**CTTTCGTGTCATCtGAGACCt |
| oCEM 055 | Golden Gate cloning | PduN-80-91-His-T88V oligo | aGGTCTCaTGACCTCGCCGTTGTCGGCATTGTAGAT**GTG**CTTTCGTGTCATCtGAGACCt |
| oCEM 056 | Golden Gate cloning | PduN-80-91-His-T88I oligo | aGGTCTCaTGACCTCGCCGTTGTCGGCATTGTAGAT**ATT**CTTTCGTGTCATCtGAGACCt |

| | | | |
|---|---|---|---|
| oCEM 057 | Golden Gate cloning | PduN-80-91-His-T88S oligo | aGGTCTCaTGACCTCGCCGTTGTCGGCATTGTAGAT**AGC**CTTTCGTGTCATCtGAGACCt |
| oCEM 058 | Golden Gate cloning | PduN-80-91-His-T88Y oligo | aGGTCTCaTGACCTCGCCGTTGTCGGCATTGTAGAT**TAT**CTTTCGTGTCATCtGAGACCt |
| oCEM 059 | Golden Gate cloning | PduN-80-91-His-T88H oligo | aGGTCTCaTGACCTCGCCGTTGTCGGCATTGTAGAT**CAT**CTTTCGTGTCATCtGAGACCt |
| oCEM 060 | Golden Gate cloning | PduN-80-91-His-T88R oligo | aGGTCTCaTGACCTCGCCGTTGTCGGCATTGTAGAT**CGC**CTTTCGTGTCATCtGAGACCt |
| oCEM 061 | Golden Gate cloning | PduN-80-91-His-T88N oligo | aGGTCTCaTGACCTCGCCGTTGTCGGCATTGTAGAT**AAC**CTTTCGTGTCATCtGAGACCt |
| oCEM 062 | Golden Gate cloning | PduN-80-91-His-T88D oligo | aGGTCTCaTGACCTCGCCGTTGTCGGCATTGTAGAT**GAT**CTTTCGTGTCATCtGAGACCt |
| oCEM 063 | Golden Gate cloning | PduN-80-91-His-T88C oligo | aGGTCTCaTGACCTCGCCGTTGTCGGCATTGTAGAT**TGC**CTTTCGTGTCATCtGAGACCt |
| oADJ 001 | QuikChange | *PduN-G52A* For | GAAGTGGCCGTGGACTCCGTCGCGGCGGGCGTCGGC |
| oADJ 002 | QuikChange | *PduN-G52A* Rev | AACCAGTTCGCCGACGCCCGCCGCGACGGAGTCCAC |
| oADJ 003 | QuikChange | *PduN-G52C* For | GAAGTGGCCGTGGACTCCGTCTGCGCGGGCGTCGGC |
| oADJ 004 | QuikChange | *PduN-G52C* Rev | AACCAGTTCGCCGACGCCCGCGCAGACGGAGTCCAC |
| oADJ 005 | QuikChange | *PduN-G52D* For | GAAGTGGCCGTGGACTCCGTCGATGCGGGCGTCGGC |
| oADJ 006 | QuikChange | *PduN-G52D* Rev | AACCAGTTCGCCGACGCCCGCATCGACGGAGTCCAC |

| | | | |
|---|---|---|---|
| oADJ 007 | QuikChange | *PduN-G52E* For | GAAGTGGCCGTGGACTCCGTC GAAGCGGGCGTCGGC |
| oADJ 008 | QuikChange | *PduN-G52E* Rev | AACCAGTTCGCCGACGCCCGC TTCGACGGAGTCCAC |
| oADJ 009 | QuikChange | *PduN-G52F* For | GAAGTGGCCGTGGACTCCGTC TTTGCGGGCGTCGGC |
| oADJ 010 | QuikChange | *PduN-G52F* Rev | AACCAGTTCGCCGACGCCCGC AAAGACGGAGTCCAC |
| oADJ 011 | QuikChange | *PduN-G52H* For | GAAGTGGCCGTGGACTCCGTC CATGCGGGCGTCGGC |
| oADJ 012 | QuikChange | *PduN-G52H* Rev | AACCAGTTCGCCGACGCCCGC ATGGACGGAGTCCAC |
| oADJ 013 | QuikChange | *PduN-G52I* For | GAAGTGGCCGTGGACTCCGTC ATTGCGGGCGTCGGC |
| oADJ 014 | QuikChange | *PduN-G52I* Rev | AACCAGTTCGCCGACGCCCGC AATGACGGAGTCCAC |
| oADJ 015 | QuikChange | *PduN-G52K* For | GAAGTGGCCGTGGACTCCGTC AAAGCGGGCGTCGGC |
| oADJ 016 | QuikChange | *PduN-G52K* Rev | AACCAGTTCGCCGACGCCCGC TTTGACGGAGTCCAC |
| oADJ 017 | QuikChange | *PduN-G52L* For | GAAGTGGCCGTGGACTCCGTC CTGGCGGGCGTCGGC |
| oADJ 018 | QuikChange | *PduN-G52L* Rev | AACCAGTTCGCCGACGCCCGC CAGGACGGAGTCCAC |
| oADJ 019 | QuikChange | *PduN-G52M* For | GAAGTGGCCGTGGACTCCGTC ATGGCGGGCGTCGGC |
| oADJ 020 | QuikChange | *PduN-G52M* Rev | AACCAGTTCGCCGACGCCCGC CATGACGGAGTCCAC |
| oADJ 021 | QuikChange | *PduN-G52N* For | GAAGTGGCCGTGGACTCCGTC AACGCGGGCGTCGGC |
| oADJ 022 | QuikChange | *PduN-G52N* Rev | AACCAGTTCGCCGACGCCCGC GTTGACGGAGTCCAC |

| | | | |
|---|---|---|---|
| oADJ 023 | QuikChange | *PduN-G52P* For | GAAGTGGCCGTGGACTCCGTC CCGGCGGGCGTCGGC |
| oADJ 024 | QuikChange | *PduN-G52P* Rev | AACCAGTTCGCCGACGCCCGC CGGGACGGAGTCCAC |
| oADJ 025 | QuikChange | *PduN-G52Q* For | GAAGTGGCCGTGGACTCCGTC CAGGCGGGCGTCGGC |
| oADJ 026 | QuikChange | *PduN-G52Q* Rev | AACCAGTTCGCCGACGCCCGC CTGGACGGAGTCCAC |
| oADJ 027 | QuikChange | *PduN-G52R* For | GAAGTGGCCGTGGACTCCGTC CGCGCGGGCGTCGGC |
| oADJ 028 | QuikChange | *PduN-G52R* Rev | AACCAGTTCGCCGACGCCCGC GCGGACGGAGTCCAC |
| oADJ 029 | QuikChange | *PduN-G52S* For | GAAGTGGCCGTGGACTCCGTC AGCGCGGGCGTCGGC |
| oADJ 030 | QuikChange | *PduN-G52S* Rev | AACCAGTTCGCCGACGCCCGC GCTGACGGAGTCCAC |
| oADJ 031 | QuikChange | *PduN-G52T* For | GAAGTGGCCGTGGACTCCGTC ACCGCGGGCGTCGGC |
| oADJ 032 | QuikChange | *PduN-G52T* Rev | AACCAGTTCGCCGACGCCCGC GGTGACGGAGTCCAC |
| oADJ 033 | QuikChange | *PduN-G52V* For | GAAGTGGCCGTGGACTCCGTC GTGGCGGGCGTCGGC |
| oADJ 034 | QuikChange | *PduN-G52V* Rev | AACCAGTTCGCCGACGCCCGC CACGACGGAGTCCAC |
| oADJ 035 | QuikChange | *PduN-G52W* For | GAAGTGGCCGTGGACTCCGTC TGGGCGGGCGTCGGC |
| oADJ 036 | QuikChange | *PduN-G52W* Rev | AACCAGTTCGCCGACGCCCGC CCAGACGGAGTCCAC |
| oADJ 037 | QuikChange | *PduN-G52Y* For | GAAGTGGCCGTGGACTCCGTC TATGCGGGCGTCGGC |
| oADJ 038 | QuikChange | *PduN-G52Y* Rev | AACCAGTTCGCCGACGCCCGC ATAGACGGAGTCCAC |

| | | | |
|---|---|---|---|
| oADJ 039 | Sequencing | Amplifies upstream of encoding region of pBAD33t For | ATGCCATAGCATTTTTATCC |
| oADJ 040 | Sequencing | Amplifies downstream of encoding region of pBAD33t Rev | GATTTAATCTGTATCAGG |
| TMD P021 | Sequencing, recombineering | Amplify from upstream of *pduD* locus For | ggcaacaggttatcgcctgc |
| TMD P022 | Sequencing, recombineering | Amplify from downstream of *pduD* locus Rev | ccctgcaggctgttcatgc |
| TMD P072 | Sequencing | Amplify from upstream of *pduD* locus For | catccagaaagccaagctaacc |
| TMD P073 | Sequencing | Amplify from downstream of *pduD* locus Rev | cgtccagcgttttattggtgg |
| TMD P016 | Recombineering | Amplify ssD-GFPmut2 with homology upstream of *pduD* For | TTGATCCCAACGAGATTGATTAAGGGGTGAGAAatggaaattaatgaaaaattgctgcgc |
| TMD P019 | Recombineering | Amplify ssD-GFPmut2 with homology downstream of *pduD* Rev | ATTGCGTCGGTATTCATGGAGTTATCCTTTAttatttgtatagttcatccatgccatgtg |

**Supplemental Table S4.** Parameters used in systems-level kinetic model for calculations of metabolite profiles over time when the Pdu pathway is encapsulated in MCPs or MTs

| Parameter Name | MCP Case | MT Case, surface area same as MCP | MT Case, volume same as MCP | Unit |
|---|---|---|---|---|
| CDE_con | 0.462 | 0.863 | 0.462 | mM |
| CDE_tot [12] | 6000 | 6000 | 6000 | enzymes/cell |
| cell_length | 2.47E-06 | 2.47E-06 | 2.47E-06 | m |
| cell_radius | 3.75E-07 | 3.75E-07 | 3.75E-07 | m |
| cell_surface_area | 5.82E-12 | 5.82E-12 | 5.82E-12 | m^2 |
| cell_volume | 9.81E-19 | 9.81E-19 | 9.81E-19 | m^3 |
| external_volume | 5.00E-05 | 5.00E-05 | 5.00E-05 | m^3 |
| kcatCDE [4] | 300 | 300 | 300 | 1/s |
| kcatL | 100 | 100 | 100 | 1/s |
| kcatPf [5] | 55 | 55 | 55 | 1/s |
| kcatPr [5] | 6 | 6 | 6 | 1/s |
| kcatQf [6] | 55 | 55 | 55 | 1/s |
| kcatQr [6] | 6 | 6 | 6 | 1/s |
| L_con [13] | 0.1 | 0.1 | 0.1 | mM |
| mcp_surface_area | 6.16E-14 | 3.88E-13 | 3.88E-13 | m^2/tube |
| mcp_volume | 1.44E-21 | 4.85E-21 | 4.85E-21 | m^3/tube |
| Navogadro | 6.02E+23 | 6.02E+23 | 6.02E+23 | molecules/mole |
| Nmcp | 15 | 2.38 | 4.44 | MTs per cell |
| P_con | 0.694 | 1.295 | 0.694 | mM |
| P_tot [12] | 9000 | 9000 | 9000 | enzymes/cell |
| PermMCPNonPolar | 3.98E-08 | 3.98E-08 | 3.98E-08 | m/s |
| PermMCPPolar | 3.98E-08 | 3.98E-08 | 3.98E-08 | m/s |
| Q_con | 0.520 | 0.971 | 0.520 | mM |
| Q_tot [12] | 6750 | 6750 | 6750 | enzymes/cell |
| radius_mcp [14] | 7.00E-08 | 2.50E-08 | 2.50E-08 | m |
| VmaxCDEf | 138.74 | 258.98 | 138.74 | mM/s |
| VmaxLf | 10.00 | 10.00 | 10.00 | mM/s |
| VmaxPf | 38.15 | 71.22 | 38.15 | mM/s |
| VmaxPr | 4.16 | 7.77 | 4.16 | mM/s |
| VmaxQf | 28.62 | 53.41 | 28.62 | mM/s |
| VmaxQr | 3.12 | 5.83 | 3.12 | mM/s |
| KmCDEPropanediol [4] | 0.5 | 0.5 | 0.5 | mM |
| KmPfPropionaldehyde [5] | 15 | 15 | 15 | mM |
| KmPrPropionyl [5] | 95 | 95 | 95 | mM |
| KmQfPropionaldehyde [6] | 15 | 15 | 15 | mM |
| KmQrPropanol [6] | 95 | 95 | 95 | mM |

| | | | | |
|---|---|---|---|---|
| KmLPropionyl | 20 | 20 | 20 | mM |
| PermCellPropanediol [15] [16] [17] | 1.00E-04 | 1.00E-04 | 1.00E-04 | m/s |
| PermCellPropionaldehyde [15] [16] [17] | 1.00E-02 | 1.00E-02 | 1.00E-02 | m/s |
| PermCellPropanol [15] [16] [17] | 1.00E-04 | 1.00E-04 | 1.00E-04 | m/s |
| PermCellPropionyl [15] [16] [17] | 1.00E-05 | 1.00E-05 | 1.00E-05 | m/s |
| PermCellPropionate[15] [16] [17] | 1.00E-07 | 1.00E-07 | 1.00E-07 | m/s |

**Supplemental Figure S1.** Transmission electron micrographs of thin cell sections of *S. enterica* LT2 expressing the *pdu* operon in (a) a PduN knockout strain and (b) a wild type strain. Arrows indicate a cell division event. In the PduN knockout strain (a), Pdu MTs can be seen blocking cell division, whereas Pdu MCPs in the wild type strain (b) do not cause cell division defects.

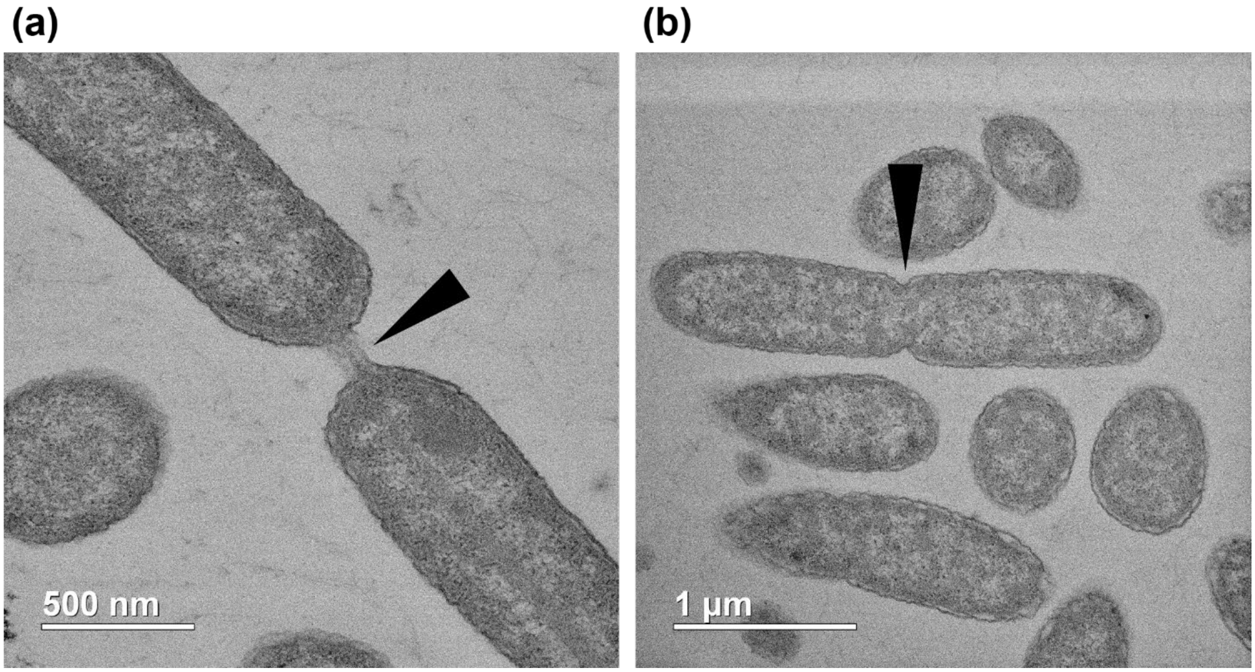

**Supplemental Figure S2.** Additional transmission electron micrographs on Pdu MTs.

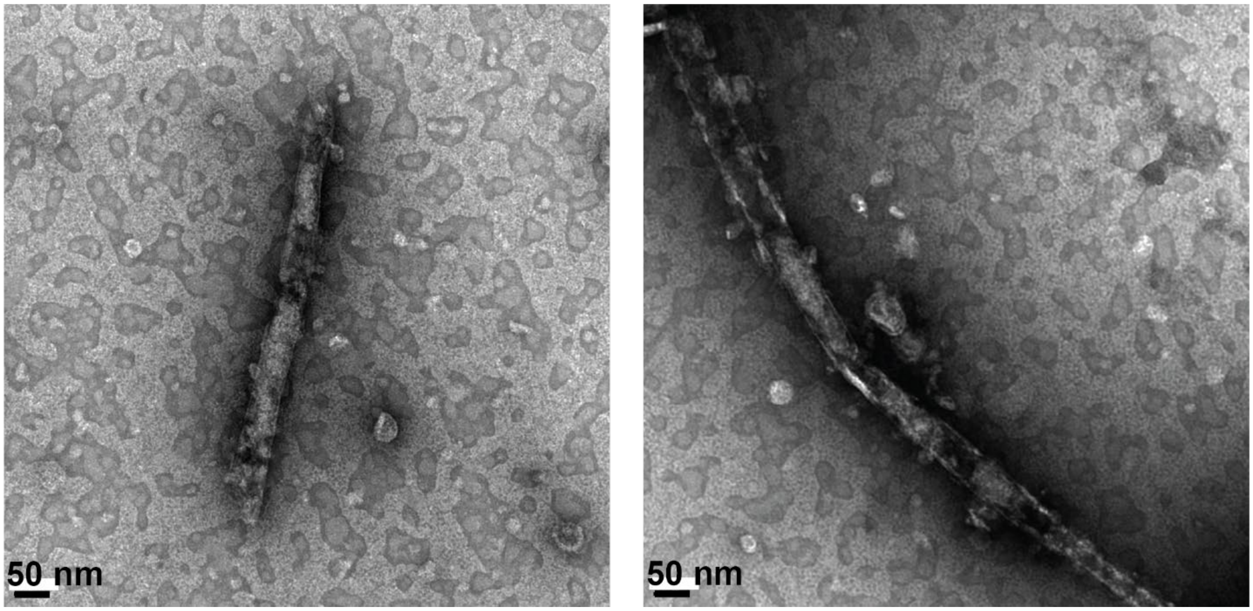

**Supplemental Figure S3.** Change in percentage of cells containing elongated, Pdu microtube (MT) structures as induction of PduN-FLAG off a plasmid is increased in a *pduN* knockout strain (ΔPduN). MT formation was visualized using a genomically integrated fluorescent reporter (ssD-GFP, integrated at the *pduD* locus [18]). WT indicates the wild type strain, and bars labeled with percentages are the PduN knockout strain with a plasmid encoding for FLAG-tagged PduN under an arabinose inducible promoter (pBAD33t-PduN-FLAG). Percentages indicate final arabinose concentration (w/w) in the culture.

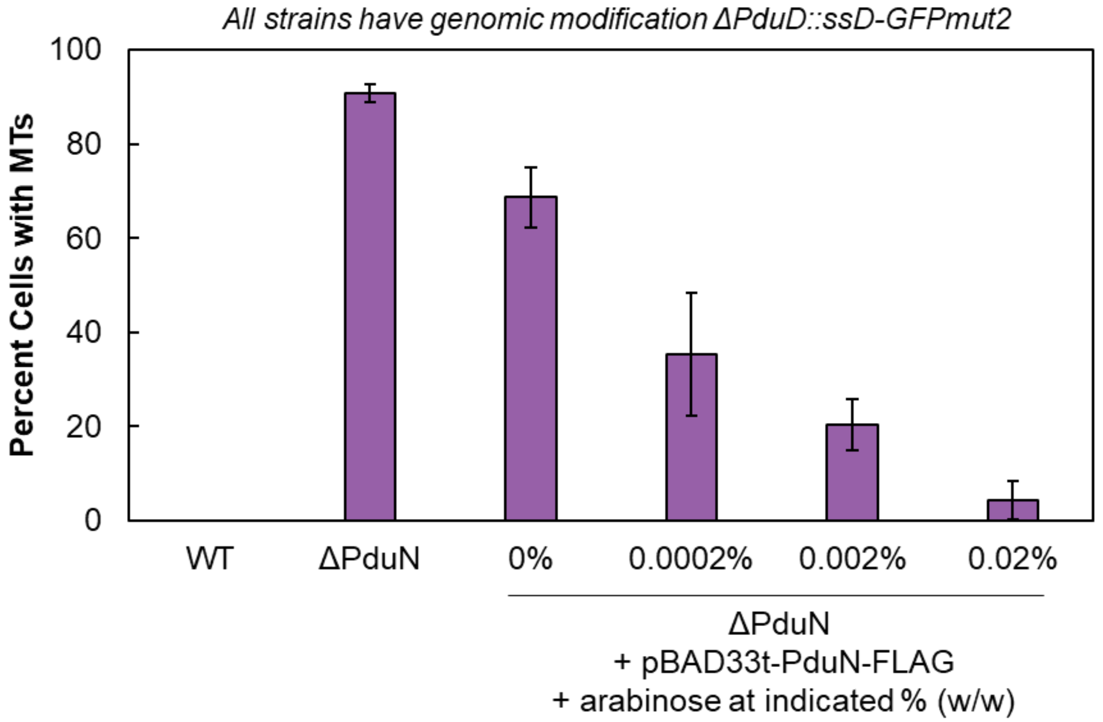

**Supplemental Figure S4.** Tornado plot showing changes in the maximum external propionaldehyde level in the MCP model with 10% increases and decreases in specified model parameters.

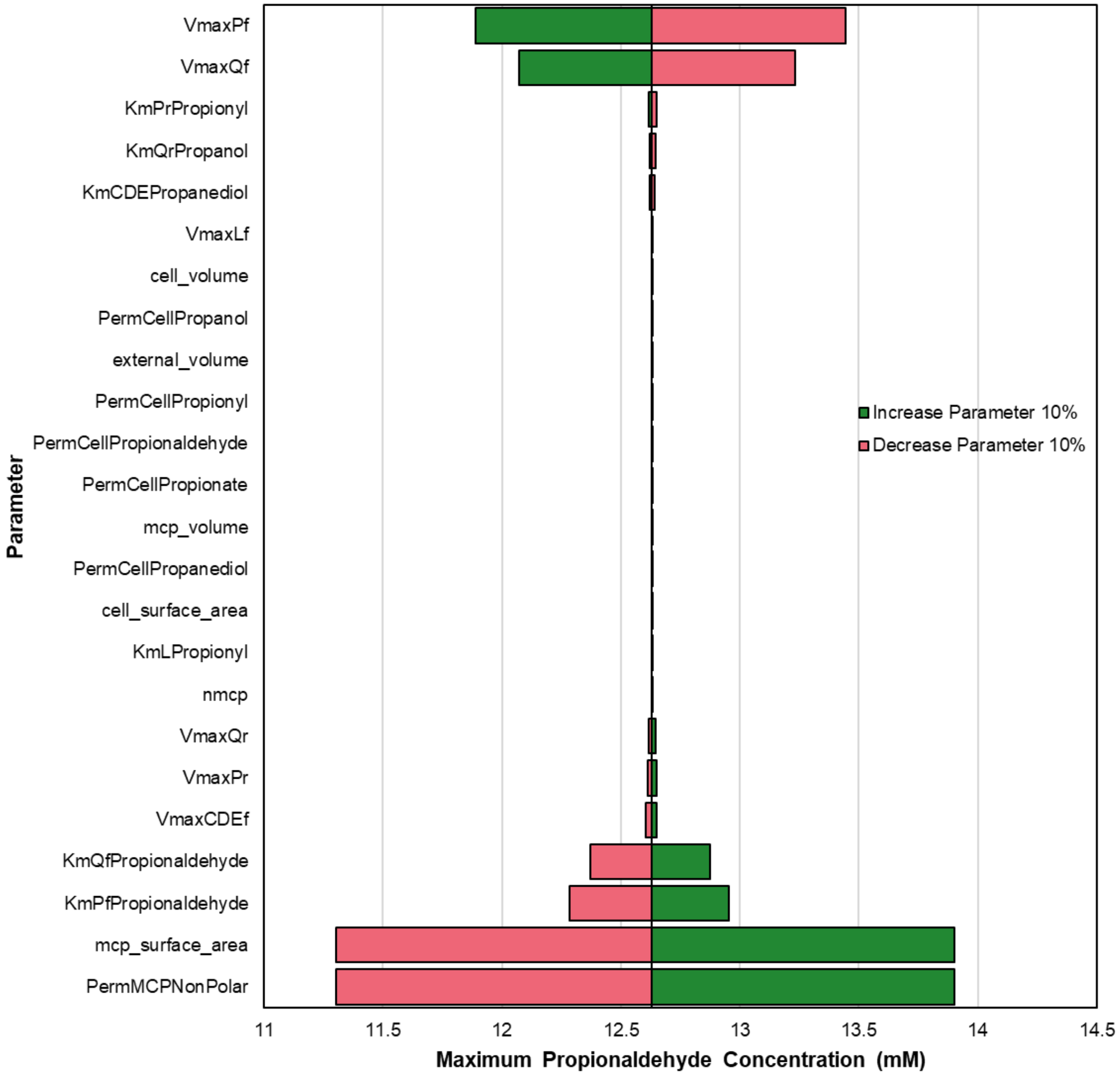

**Supplemental Figure S5.** Tornado plot showing changes in the maximum external propionaldehyde level in the microtube (MT) model (total MT volume equal to total MCP volume; enzyme concentration same as in MCP model; 190% increase in surface area compared to MCP surface area) with 10% increases and decreases in specified model parameters.

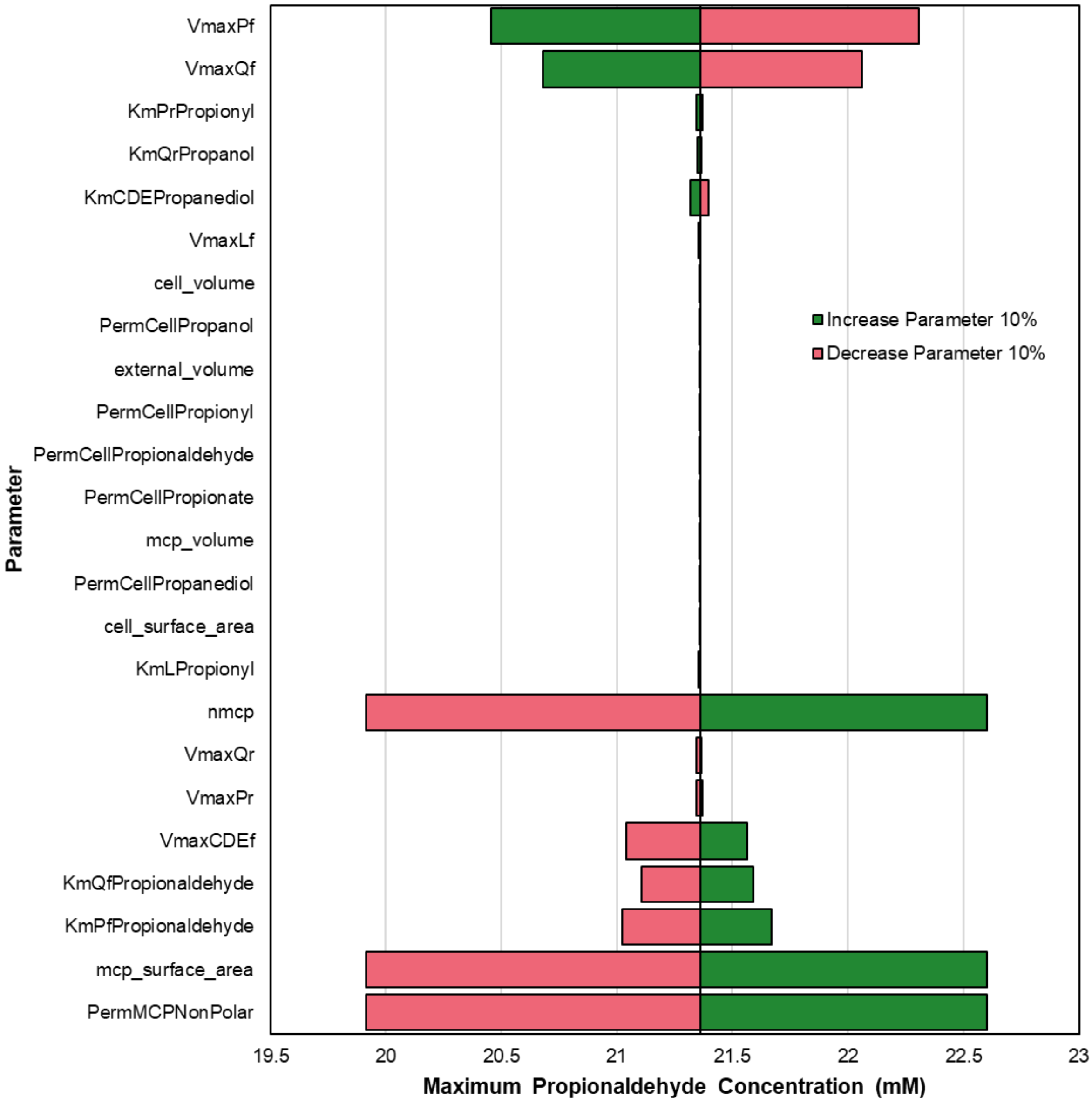

**Supplemental Figure S6.** Mapping between bending angle, $\theta_B$ (degrees), and the distance between the centers of mass of the pentamer and hexamer, $z$ (nm).

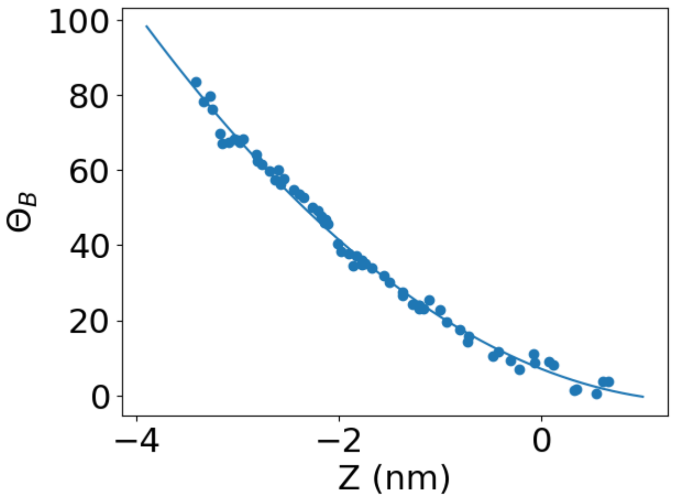

**Supplemental Figure S7.** Histograms showing the parallel "windows" used to calculate the potential of mean force. The overlap of the windows is shown by the distance between the centers of mass of the pentamer and hexamer, *z* (nm).

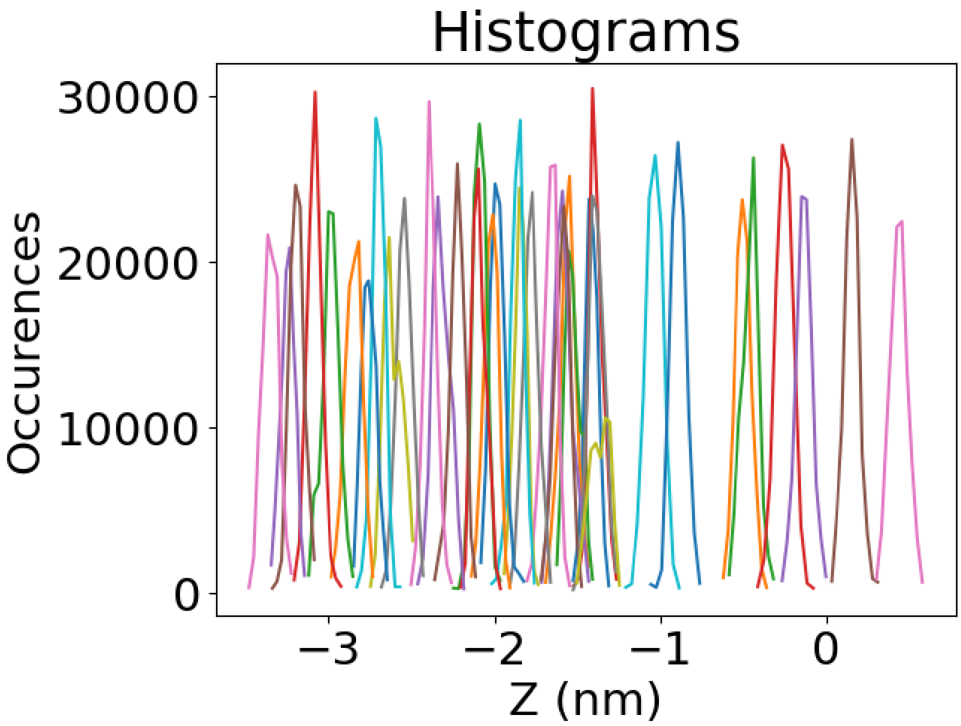

### Bibliography

[1] S. Páll, M.J. Abraham, C. Kutzner, B. Hess, and E. Lindahl, "Tackling exascale software challenges in molecular dynamics simulations with GROMACS"; pp. 3–27 in *Solving software challenges for exascale*. Edited by S. Markidis and E. Laure. Springer International Publishing Switzerland, London, 2015.

[2] E.M. Sampson, T.A. Bobik, Microcompartments for B12-Dependent 1,2-Propanediol Degradation Provide Protection from DNA and Cellular Damage by a Reactive Metabolic Intermediate, J. Bacteriol. 190 (2008) 2966-2971. https://doi.org/10.1128/JB.01925-07.

[3] C.M. Jakobson, D. Tullman-Ercek, M.F. Slininger, N.M. Mangan, A systems-level model reveals that 1,2-Propanediol utilization microcompartments enhance pathway flux through intermediate sequestration, PLoS Comput. Biol. 13 (2017) e1005525. https://doi.org/10.1371/journal.pcbi.1005525.

[4] W.W. Bachovichin, R.G. Eagar, Jr., K.W. Moore, J.H. Richards, Mechanism of action of adenosylcobalamin: glycerol and other substrate analogs as substrates and inactivators for propanediol dehydratase - kinetics, stereospecificity, and mechanism, Biochemistry. 16 (1977) 1082-1092. https://doi.org/10.1021/bi00625a009.

[5] N.A. Leal, G.D. Havemann, T.A. Bobik, PduP is a coenzyme-a-acylating propionaldehyde dehydrogenase associated with the polyhedral bodies involved in B12-dependent 1,2-propanediol degradation by Salmonella enterica serovar Typhimurium LT2, Arch Microbiol. 180 (2003) 353-361. https://doi.org/10.1007/s00203-003-0601-0.

[6] S. Cheng, C. Fan, S. Sinha, T.A. Bobik, The PduQ Enzyme Is an Alcohol Dehydrogenase Used to Recycle $NAD^+$ Internally within the Pdu Microcompartment of *Salmonella enterica*, PLoS ONE. 7 (2012) e47144. https://doi.org/10.1371/journal.pone.0047144.

[7] M. Sutter, B. Greber, C. Aussignargues, C.A. Kerfeld, Assembly principles and structure of a 6.5-MDa bacterial microcompartment shell, Science. 356 (2017) 1293-1297. https://doi.org/10.1126/science.aan3289.

[8] B.J. Greber, M. Sutter, C.A. Kerfeld, The Plasticity of Molecular Interactions Governs Bacterial Microcompartment Shell Assembly, Structure. 27 (2019) 749-763. https://doi.org/10.1016/j.str.2019.01.017.

[9] S.A. Klein, A. Majumdar, D. Barrick, A Second Backbone: The Contribution of a Buried Asparagine Ladder to the Global and Local Stability of a Leucine-Rich Repeat Protein, Biochemistry. 58 (2019) 3480-3493. https://doi.org/10.1021/acs.biochem.9b00355.

[10] L.C. Thomason, J.A. Sawitzke, X. Li, N. Costantino, D.L. Court, Recombineering: genetic engineering in bacteria using homologous recombination, Curr. Protoc. Mol. Biol. 106 (2014) 1.16.1-1.16.39. https://doi.org/10.1002/0471142727.mb0116s106.

[11] S. Datta, N. Costantino, D.L. Court, A set of recombineering plasmids for gram-negative bacteria, Gene. 379 (2006) 109–115. https://doi.org/10.1016/j.gene.2006.04.018.

[12] M. Yang, D.M. Simpson, N. Wenner, P. Brownridge, V. M. Harman, J.C.D. Hinton, R.J. Beynon, L. Liu, Decoding the stoichiometric composition and organisation of bacterial metabolosomes, Nat. Comm. 11 (2020) 1976. https://doi.org/10.1038/s41467-020-15888-4.

[13] K.R. Able, M.H. Butler, B.E.Wright, Cellular Concentrations of Enzymes and Their Substrates, J. Theor. Biol. 143 (1990) 163-195.

[14] N.W. Kennedy, J.M. Hershewe, T.M. Nichols, E.W. Roth, C.D. Wilke, C.E. Mills, M.C. Jewett, D. Tullman-Ercek, Apparent size and morphology of bacterial microcompartments varies with technique, PLos ONE. 15 (2019) e0226395. https://doi.org/10.1371/journal.pone.0226395.

[15] E. Orbach, A. Finkelstein, The nonelectrolyte permeability of planar lipid bilayer membranes, J. Gen. Physiol. 75 (1980) 427-436.

[16] M.H. Abraham, G.S. Whiting, R. Fuchs, E.J. Chambers, Thermodynamics of solute transfer from water to hexadecane, J. Chem. Soc., Perkin Trans. 2. (1990) 291-300. https://doi.org/10.1039/P29900000291.

[17] M.M. Schantz, D.E. Martire, Determination of hydrocarbon-water partition coefficients from chromatographic data and based on solution thermodynamics and theory, J. Chromatogr. A. 391 (1987) 35-51.

[18] T.M. Nichols, N.W. Kennedy, D. Tullman-Ercek, A genomic integration platform for heterologous cargo encapsulation in 1,2-propanediol utilization bacterial microcompartments, Biochem. Eng. J. 156 (2020) 107496. https://doi.org/10.1016/j.bej.2020.107496.
