## Supplemental Document 2 for "Vertex protein PduN tunes encapsulated pathway performance by dictating bacterial metabolosome morphology"

#### 1 Introduction

The kinetic, systems-level model for the 1,2-propanediol utilization (Pdu) pathway used in this manuscript was developed based on the original work of Jakobson *et al.*,[1] with some modifications.
Specific modifications include:

- The assumption that the cytosol of the cell is well-mixed.
- The explicit modeling of the metabolite concentrations in the external media.
- The incorporation of cell growth over time.
- The addition of reaction terms for the conversion of propionyl-CoA into propionate in the cytosol by PduL/PduW.
- The consideration of reverse reaction rates for PduP and PduQ enzymes.

Compared to the previously published work, where analysis focused on the steady-state condition, we focus the analysis in this manuscript on the change in metabolite profiles over time in a batch reactor, as these conditions match those used to generate experimental data.

#### 2 Model Assumptions

We make the following assumptions in our model:

1. At time $t$, there are $N(t)$ identical, non-interacting cells in a well mixed solution.
2. The substrates 1,2-propanediol, propionaldehyde, propionyl-CoA, propionate, and 1-propanol passively diffuse across the cell membrane at rates specified by permeability parameters.
3. The substrates 1,2-propanediol, propionaldehyde, propionyl-CoA, propionate, and 1-propanol passively diffuse across the microcompartment (MCP) shell over the entire surface of the spherical MCP at rates specified by permeability parameters.
4. The substrates 1,2-propanediol, propionaldehyde, propionyl-CoA, propionate, and 1-propanol passively diffuse across the microtube (MT) shell along the long axis of the cylindrical MT at rates specified by permeability parameters.
5. There are $n_{\text{MCP}}$ non-interacting MCPs or MTs in the cytosol of each cell.
6. Reactions catalyzed by PduCDE, PduP, and PduQ, forward or reverse, can only occur in the MCP or MT

7. Reactions catalyzed by PduL/W can only occur in the cytosol.
8. The external media, cytosol of the cell, and internal MCP/MT volume are well-mixed such that the concentration in each compartment is assumed uniform at any given point in time.
9. The volume of the external media up to leading order is the volume of the entire culture.
10. The volume of the cytosol up to leading order is the volume of the cell.
11. All enzymes behave according to Michaelis-Menten kinetics.

### 3 Chemical Reactions

The following reactions are considered in our model, with the assumptions described above:

$$\text{1,2-propanediol} \xrightarrow{\text{PduCDE}} \text{Propionaldehyde}$$

$$\text{propionaldehyde} \xleftrightarrow{\text{PduQ}} \text{1-propanol}$$

$$\text{propionaldehyde} \xleftrightarrow{\text{PduP}} \text{propionyl-CoA}$$

$$\text{propionyl-CoA+} \xrightarrow{\text{PduLW}} \text{propionate}$$

### 4 Equations Used In Mathematical Model

The differential equations described below were integrated forward in time from a starting condition of 55 mM 1,2-propanediol in the external media, the starting condition for our growth curve.

#### 4.1 Variable Definition

Concentrations of all substrates are defined by the following variables:

- $P_i$: 1,2-propanediol concentration in volume $i$ (MCP/MT, cytosol, or external media)
- $A_i$: propionaldehyde concentration in volume $i$ (MCP/MT, cytosol, or external media)
- $Pol_i$: 1-propanol concentration in volume $i$ (MCP/MT, cytosol, or external media)
- $PCoA_i$: propionyl-CoA concentration in volume $i$ (MCP/MT, cytosol, or external media)
- $Pate_i$: propionate concentration in volume $i$ (MCP/MT, cytosol, or external media)

Other constants and variables used in the model are defined follows:

- $V_{\text{compartment}}$: volume of the compartment (MCP or MT)
- $SA_{\text{compartment}}$: surface area of the compartment (MCP or MT); for the MT, this only includes the long axis of the cylinder
- $Vol_{\text{cell}}$: volume of the cell
- $SA_{\text{cell}}$: surface area of the cell
- $Perm_i^j$: permeability of substrate $i$ at interface $j$ (cell surface or MCP/MT surface)
- $R_i(X_j)$: reaction rate of enzyme $i$ as a function of concentrations $X$ in a given volume $j$ (MCP/MT, cytosol, or external media)
- $K_{\text{M}}^{i,j}$: Michaelis constant of enzyme $i$ for substrate $j$

- $V_{\max}^{i,j}$: Maximum reaction velocity of enzyme $i$ for substrate $j$
- $N(t)$: Number of cells at time $t$, calculated from the experimental growth profile.
- MT: Refers to MT
- MCP: Refers to MCP
- $n_{\text{MCP}}$: Number of MCPs/MTs per cell

#### 4.2 Differential Equations: MCP or MT

The differential equations for the MCP or MT volume were as follows:

$$
\begin{aligned}
\frac{dP_{\text{MCP/MT}}}{dt} =& - R_{\text{PduCDE}}\left(X_{\text{MCP/MT}}\right) + \frac{Perm^{\text{P}}_{\text{MCP/MT}} SA_{\text{MCP/MT}}}{Vol_{\text{MCP/MT}}}\left(P_{\text{cytosol}} - P_{\text{MCP/MT}}\right) \\
\frac{dA_{\text{MCP/MT}}}{dt} =& R_{\text{PduCDE}}\left(X_{\text{MCP/MT}}\right) - R_{\text{PduP},f}\left(X_{\text{MCP/MT}}\right) + R_{\text{PduP},r}\left(X_{\text{MCP/MT}}\right) \\
& - R_{\text{PduQ},f}\left(X_{\text{MCP/MT}}\right) + R_{\text{PduQ},r}\left(X_{\text{MCP/MT}}\right) \\
& + \frac{Perm^{\text{A}}_{\text{MCP/MT}} SA_{\text{MCP/MT}}}{Vol_{\text{MCP/MT}}}\left(A_{\text{cytosol}} - A_{\text{MCP/MT}}\right) \\
\frac{dPol_{\text{MCP/MT}}}{dt} =& R_{\text{PduQ},f}\left(X_{\text{MCP/MT}}\right) - R_{\text{PduQ},r}\left(X_{\text{MCP/MT}}\right) \\
& + \frac{Perm^{\text{Pol}}_{\text{MCP/MT}} SA_{\text{MCP/MT}}}{Vol_{\text{MCP/MT}}}\left(Pol_{\text{cytosol}} - Pol_{\text{MCP/MT}}\right) \\
\frac{dPCoA_{\text{MCP/MT}}}{dt} =& R_{\text{PduP},f}\left(X_{\text{MCP/MT}}\right) - R_{\text{PduP},r}\left(X_{\text{MCP/MT}}\right) \\
& + \frac{Perm^{\text{PCoA}}_{\text{MCP/MT}} SA_{\text{MCP/MT}}}{Vol_{\text{MCP/MT}}}\left(PCoA_{\text{cytosol}} - PCoA_{\text{MCP/MT}}\right) \\
\frac{dPate_{\text{MCP/MT}}}{dt} =& \frac{Perm^{\text{Pate}}_{\text{MCP/MT}} SA_{\text{MCP/MT}}}{Vol_{\text{MCP/MT}}}\left(Pate_{\text{cytosol}} - Pate_{\text{MCP/MT}}\right)
\end{aligned}
$$

#### 4.3 Differential Equations: Cytosol

The differential equations for the cytosol of the cell were as follows:

$$
\begin{aligned}
\frac{dP_{\text{cytosol}}}{dt} =& \frac{n_{\text{MCP}} Perm^{\text{P}}_{\text{MCP/MT}} SA_{\text{MCP/MT}}}{Vol_{\text{MCP/MT}}} \left(P_{\text{MCP/MT}} - P_{\text{cytosol}}\right) \\
&+ \frac{Perm^{\text{P}}_{\text{cell}} SA_{\text{cell}}}{Vol_{\text{cell}}} \left(P_{\text{external}} - P_{\text{cytosol}}\right) \\
\frac{dA_{\text{cytosol}}}{dt} =& \frac{n_{\text{MCP}} Perm^{\text{A}}_{\text{MCP/MT}} SA_{\text{MCP/MT}}}{Vol_{\text{MCP/MT}}} \left(A_{\text{MCP/MT}} - A_{\text{cytosol}}\right) \\
&+ \frac{Perm^{\text{A}}_{\text{cell}} SA_{\text{cell}}}{Vol_{\text{cell}}} \left(A_{\text{external}} - A_{\text{cytosol}}\right) \\
\frac{dPol_{\text{cytosol}}}{dt} =& \frac{n_{\text{MCP}} Perm^{\text{Pol}}_{\text{MCP/MT}} SA_{\text{MCP/MT}}}{Vol_{\text{MCP/MT}}} \left(Pol_{\text{MCP/MT}} - Pol_{\text{cytosol}}\right) \\
&+ \frac{Perm^{\text{Pol}}_{\text{cell}} SA_{\text{cell}}}{Vol_{\text{cell}}} \left(Pol_{\text{external}} - Pol_{\text{cytosol}}\right) \\
\frac{dPCoA_{\text{cytosol}}}{dt} =& - R_{\text{PduLW}} \left(X_{\text{cytosol}}\right) \\
&+ \frac{n_{\text{MCPB}} Perm^{\text{PCoA}}_{\text{MCP/MT}} SA_{\text{MCP/MT}}}{Vol_{\text{MCP/MT}}} \left(PCoA_{\text{MCP/MT}} - PCoA_{\text{cytosol}}\right) \\
&+ \frac{Perm^{\text{PCoA}}_{\text{cell}} SA_{\text{cell}}}{Vol_{\text{cell}}} \left(PCoA_{\text{external}} - PCoA_{\text{cytosol}}\right) \\
\frac{dPate_{\text{cytosol}}}{dt} =& R_{\text{PduLW}} \left(X_{\text{cytosol}}\right) \\
&+ \frac{n_{\text{MCP}} Perm^{\text{Pate}}_{\text{MCP/MT}} SA_{\text{MCP/MT}}}{Vol_{\text{MCP/MT}}} \left(Pate_{\text{MCP/MT}} - Pate_{\text{cytosol}}\right) \\
&+ \frac{Perm^{\text{Pate}}_{\text{cell}} SA_{\text{cell}}}{Vol_{\text{cell}}} \left(Pate_{\text{external}} - Pate_{\text{cytosol}}\right)
\end{aligned}
$$

#### 4.4 Differential Equations: External Media

The differential equations for the external media of the cell were as follows:

$$
\begin{aligned}
\frac{dP_{\text{external}}}{dt} =& N(t) \frac{Perm^{\text{P}}_{\text{cell}} SA_{\text{cell}}}{Vol_{\text{cell}}} \left(P_{\text{external}} - P_{\text{cytosol}}\right) \\
\frac{dA_{\text{external}}}{dt} =& N(t) \frac{Perm^{\text{A}}_{\text{cytosol}} SA_{\text{cell}}}{Vol_{\text{cell}}} \left(A_{\text{external}} - A_{\text{cytosol}}\right) \\
\frac{dPol_{\text{external}}}{dt} =& N(t) \frac{Perm^{\text{Pol}}_{\text{cytosol}} SA_{\text{cell}}}{Vol_{\text{cell}}} \left(Pol_{\text{external}} - Pol_{\text{cytosol}}\right) \\
\frac{dPCoA_{\text{external}}}{dt} =& N(t) \frac{Perm^{\text{PCoA}}_{\text{cytosol}} SA_{\text{cell}}}{Vol_{\text{cell}}} \left(PCoA_{\text{external}} - PCoA_{\text{cytosol}}\right) \\
\frac{dPate_{\text{external}}}{dt} =& N(t) \frac{Perm^{\text{Pate}}_{\text{cytosol}} SA_{\text{cell}}}{Vol_{\text{cell}}} \left(Pate_{\text{external}} - Pate_{\text{cytosol}}\right)
\end{aligned}
$$

#### 4.5 Definition of Reaction Rate

Reaction rates were assumed to follow Michaelis-Menten kinetics

$$R_{\mathrm{PduCDE}} = V_{\mathrm{max}}^{\mathrm{PduCDE}} \frac{P_{\mathrm{MCP/MT}}}{K_{\mathrm{M}}^{\mathrm{PduCDE}} + P_{\mathrm{MCP/MT}}}$$

$$R_{\mathrm{PduP},f} = V_{\mathrm{max}}^{\mathrm{PduP},f} \frac{A_{\mathrm{MCP/MT}}}{K_{\mathrm{M}}^{\mathrm{PduP,f}} + A_{\mathrm{MCP/MT}}}$$

$$R_{\mathrm{PduP},r} = V_{\mathrm{max}}^{\mathrm{PduP},r} \frac{PCoA_{\mathrm{MCP/MT}}}{K_{\mathrm{M}}^{\mathrm{PduP,r}} + PCoA_{\mathrm{MCP/MT}}}$$

$$R_{\mathrm{PduQ},f} = V_{\mathrm{max}}^{\mathrm{PduQ},f} \frac{A_{\mathrm{MCP/MT}}}{K_{\mathrm{M}}^{\mathrm{PduQ,f}} + A_{\mathrm{MCP/MT}}}$$

$$R_{\mathrm{PduQ},r} = V_{\mathrm{max}}^{\mathrm{PduQ},r} \frac{Pol_{\mathrm{MCP/MT}}}{K_{\mathrm{M}}^{\mathrm{PduQ,r}} + Pol_{\mathrm{MCP/MT}}}$$

$$R_{\mathrm{PduLW}} = V_{\mathrm{max}}^{\mathrm{PduLW}} \frac{PCoA_{\mathrm{cytosol}}}{K_{\mathrm{M}}^{\mathrm{PduLW}} + PCoA_{\mathrm{cytosol}}}$$

### References

[1] Christopher M. Jakobson, Danielle Tullman-Ercek, Marilyn F. Slininger, and Niall M. Mangan. A systems-level model reveals that 1,2-propanediol utilization microcompartments enhance pathway flux through intermediate sequestration. *PLOS Computational Biology*, 13(5):1–24, 05 2017.
